# gAIRR-wgs: An Algorithm to Genotype T Cell Receptor Alleles Using Whole Genome Sequencing Data

**DOI:** 10.1101/2025.10.11.681280

**Authors:** Kuan-Ta Huang, Yu-Hsuan Yang, Mao-Jan Lin, Sheng-Kai Lai, Ting-Hsuan Chou, Chieh-Yu Lee, Tsung-Kai Hung, Chia-Lang Hsu, Ya-Chien Yang, Chien-Yu Chen, Pei-Lung Chen, Jacob Shu-Jui Hsu

## Abstract

*T cell receptor* (*TR*) genes, including variable (TR_V), diversity (TR_D), and joining (TR_J) segments, exhibit allelic diversity that is critical to adaptive immunity. Growing evidence has identified associations between *TR* genes and immune-related diseases. Germline variants may influence *TR* gene function and subsequent usage, highlighting the importance of accurate *TR* allele profiling. However, accurately identifying germline *TR* from standard WGS data remains challenging due to short read lengths, limited depth, and high sequence similarity. To address these challenges, we developed gAIRR-wgs, for WGS-based *TR* allele typing. By incorporating novel alleles from HPRC individuals, gAIRR-wgs exhibited excellent performance in allele calling, with F1 scores of 100.0% for TR_D, 99.8% for TR_J, and 98.3% for TR_V. Applying this pipeline to 1,492 individuals from the Taiwan Biobank (TWB), we identified 449 novel *TR* alleles, 277 of which overlapped with HPRC release 1 data of mixed ethnicity and are absent in the IMGT database. Further population comparison analysis revealed significant *TR* allele distribution differences across global populations, showing population-specific patterns and diversity variations between ethnic groups. We also discovered TWB-specific deletion polymorphisms affecting contiguous *TRGV* and *TRBV* genes, which are not recorded in the gnomAD database and undetected by standard structural variant callers, highlighting the need for tailored approaches to resolve complex immune gene regions. In conclusion, gAIRR-wgs enables accurate *TR* allele calling from standard WGS data using feasible computational resources and reveals substantial immunogenetic diversity in population cohorts.

## INTRODUCTION

T cell receptors, together with immunoglobulins which include both membrane-bound B cell receptors and soluble antibodies, form the adaptive immune receptor repertoires (AIRR), the foundation of adaptive immunity. T cell receptors enable T cells to recognize antigens presented by human leukocyte antigens (HLAs), driving cell-mediated immune responses. T cell receptors consist of α:β or γ:δ peptide chains, encoded by *T cell receptor* (*TR*) genes, including *TRA*, *TRB*, *TRG*, and *TRD*, which are located at distinct genomic loci: *TRA* and *TRD* are co-located on chromosome 14, *TRB* and *TRG* reside on chromosome 7, and an additional orphon *TRB* locus is located on chromosome 9. Each *TR* locus is composed of multiple gene segments: variable (V), diversity (D), and joining (J), which undergo somatic recombination to form functional *TR* genes during T cell development in the thymus. All *TR* loci contain V and J gene segments, while *TRB* and *TRD* additionally possess D gene segments. Within each locus, multiple copies of V, D, and J genes are arranged either in clusters or interspersed along the chromosome, and each individual gene can exist as multiple allelic variants. The germline *TR* (g*TR*) repertoire refers to the set of unrearranged V, D, and J gene segments that collectively generate *TR* diversity through recombination. The international ImMunoGeneTics information system (IMGT) database serves as the nomenclature authority and central repository for g*TR* genes, providing curated reference sequences based primarily on core coding regions (Lane et al., 2010). In this study, we classify g*TR* genes by segment type, given that genes within each segment category share similar lengths and intrinsic sequence complexity: TR_V refers to all variable genes across all *TR* loci, while TR_D and TR_J represent the diversity and joining genes, respectively. The functional diversity arising from g*TR* recombination manifests as the expressed adaptive immune receptor repertoire (exprAIRR), which has been widely associated with disease susceptibility, clinical progression, and prognosis. Non-random selected g*TR* gene combinations, have been observed across a wide range of diseases, including autoimmune diseases (Z. Chen et al., 2021; Fichtner et al., 2020; Jacobsen et al., 2017; Jia et al., 2022; Mitchell & Michels, 2020; Turcinov et al., 2023; Wang et al., 2016; Zheng et al., 2021), infectious diseases (Dolton et al., 2022; Frimpong et al., 2022; Gras et al., 2010; Li et al., 2021; Marín-Benesiu et al., 2024; Mazouz et al., 2021; Miles et al., 2011; Minervina et al., 2021; Shao et al., 2022; Shomuradova et al., 2020), severe cutaneous adverse reactions (SCARs) (Pan et al., 2019), and various cancers (Y. T. Chen et al., 2021; Huda et al., 2021; Keane et al., 2017; Wang et al., 2016; Yu et al., 2024; Zhang et al., 2024). Despite growing evidence for the role of exprAIRR in diseases, the impact of g*TR* allele variation remains insufficiently studied. Distinct g*TR* allelic forms of the same gene can lead to altered V(D)J recombination outcomes, including the production of nonfunctional transcripts or pseudogenes, thereby directly affecting gene availability, contributing to non-random gene usage, and ultimately shaping immune responses and disease susceptibility (Charmley et al., 1993; Omer et al., 2022). Current approaches for g*TR* allele identification can be broadly categorized into two methodologies: inference from rearranged transcripts (exprAIRR) and direct calling from germline DNA sequences. While transcript-based methods capture expressed repertoires, they cannot comprehensively assess the complete germline allelic diversity, as not all alleles are necessarily expressed or detectable in a given sample. Direct germline calling from whole-genome sequencing (WGS) data offers a more complete view of g*TR* variation; however, the high sequence similarity among *TR* gene segments poses substantial challenges for accurate allele identification from short-read technologies. Existing germline *TR*-calling tools, such as gAIRR-Suite, have demonstrated promising performance but are primarily optimized for targeted panel sequencing with ultra-high depth (>500X), making them incompatible with standard WGS data (∼30X) (Lin et al., 2022). This limitation is particularly significant given that biobanks offer an unprecedented opportunity to explore population-level g*TR* diversity, with most containing individual short-read WGS data at moderate sequencing depth (∼30X) (Bick et al., 2024; Elfatih et al., 2024; Feng et al., 2022; Flanagan et al., 2024; Halldorsson et al., 2022; Hsu et al., 2024; Pajuste & Remm, 2023). While individual-level g*TR* variants impact immune function, population-level variation provides essential biological context— not only establishing the baseline diversity against which individual deviations are interpreted, but also revealing patterns shaped by evolutionary selection and population-specific immune pressures. However, no existing tools can effectively and accurately identify *TR* alleles from the standard-depth short-read WGS data commonly available in biobanks. To address this critical gap, we developed gAIRR-wgs, a WGS-compatible module for gAIRR-Suite (Lin et al., 2022), to bring high-accuracy *TR* allele identification to standard-depth biobank data.

## RESULTS

### 1. Development of the gAIRR-wgs pipeline for *TR* genotyping using WGS data

The main challenges in calling g*TR* alleles from WGS data included lower read depth compared to targeted sequencing, shorter read lengths (150 bp), and the limited ability to leverage paired-end information due to the g*TR* structure (**Supplementary Figure 1**). Additionally, WGS data encompasses the entire genome, introducing substantial computational challenges and background noise from non-target regions. To address these limitations, we developed gAIRR-wgs, a specialized module of gAIRR-suite optimized for calling *TR* alleles from WGS data (**Figure 1**).

**Figure 1.**
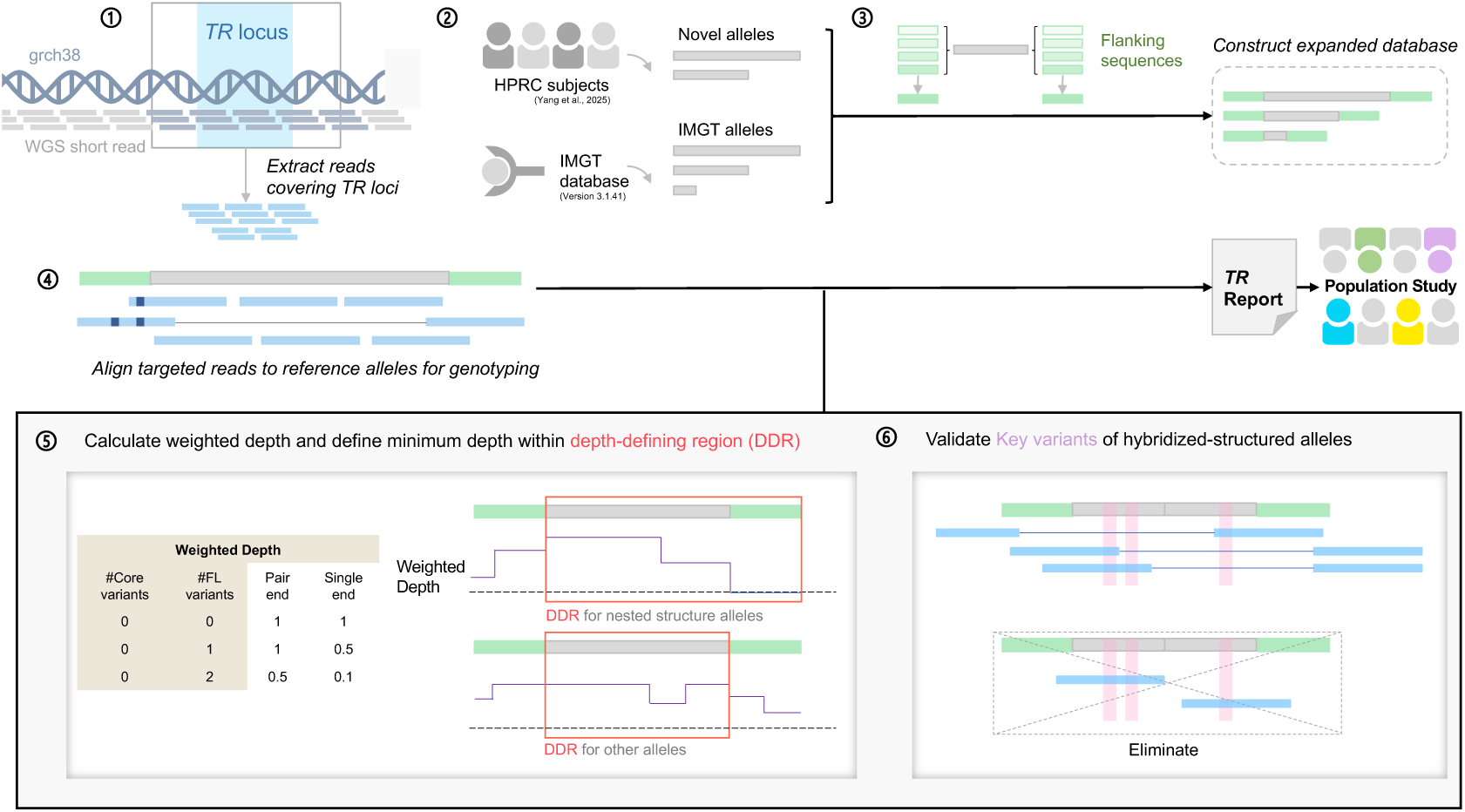
Overview of the gAIRR-wgs pipeline for *TR* allele calling from WGS data. *TR*-related reads were extracted from WGS data (top left; ➀), and the allele reference database was expanded by incorporating novel alleles from 47 HPRC subjects (top; ➁) and appending consensus-based flanking sequences (top right; ➂). Reads were then aligned to the expanded database to identify both candidate novel alleles and IMGT-documented alleles (top; ➃). Allele support was quantified using a weighted depth model that integrates both single-end and paired-end reads, with depths calculated either within customized depth-defining regions (DDR) for each allele (bottom left; ➄). Hybridized-structured alleles were further validated by requiring proper paired reads to support all key distinguishing variants (bottom right; ➅).

To optimize computational efficiency and reduce background noise, we implemented a targeted read extraction strategy. WGS reads were first aligned to the GRCh38 reference genome using BWA-MEM, followed by extraction of reads mapping to extended *TR* genomic regions using SAMtools (**Supplementary Figure 2**). Rather than using exact *TR* gene coordinates on reference genome, we extended the target regions by approximately 2 kb to capture informative reads potentially located outside strict gene boundaries. Unmapped reads were also collected to account for potential discrepancies between the reference genome and individual-specific sequences. This extraction process dramatically improved computational performance. Prior to extraction, analysis of a single sample required 6-7 hours with maximum memory usage exceeding 500 GB. After extraction, paired-end read files were reduced in size from approximately 26 GB each to 380 MB each, enabling the same analyses to be completed in 12-15 minutes with maximum memory usage of approximately 5 GB, representing a >25-fold improvement in both time and memory efficiency.

We expanded the reference database by incorporating 335 novel *TR* alleles previously discovered and validated using long-read sequencing data (Yang et al., 2025) (**Figure 1**). To improve read mapping efficiency and allele discrimination, we extended each reference sequence with 50 bp flanking sequences on both ends (**Figure 1**, **Supplementary Figure 1**). These flanking sequences were derived from HPRC assembly data using a consensus-based strategy that included sequences supported by at least 25% of the HPRC samples, providing a balance between sequence diversity and reliability (**Figure 1**, **Supplementary Figure 3**). Since flanking sequences represented merged consensus versions that might not perfectly match individual-specific sequences, we implemented some flexibilities in subsequent algorithms that allows reads to contain variants in flanking regions while maintaining perfect matches in core gene regions during depth calculation.

Following the original gAIRR-panel workflow for targeted-sequencing, we performed initial alignment using BWA-MEM in paired-end mode to map reads to the expanded reference database (**Figure 1**). Candidate novel alleles were identified when variants were supported by at least 25% of reads at specific positions. However, due to the incorporation of flanking regions in the database, novel allele calling was restricted to variants located within core gene regions, as defined by IMGT nomenclature guidelines (Lefranc, 2011), to avoid calling flanking sequence polymorphisms as novel alleles. Candidate novel alleles were then incorporated into the database for following allele calling. To address two common sources of alignment ambiguity, including hybridized-structured alleles and nested-structured alleles, we implemented a comprehensive strategy combining alignment parameter modifications, sophisticated read weighting systems, and validation mechanisms. Hybridized-structured alleles present a particular challenge during alignment, as their 5’ region may share similarity with one allele while their 3’ region resembles another allele within the same gene (**Supplementary Table 1**). During alignment, reads derived from other alleles may be misaligned to these alleles with hybridized-structured, creating artificial support and leading to false positive calls. This issue is particularly pronounced in WGS data with 150 bp reads and lower sequencing depth (30x), where alignment ambiguity is more severe. To mitigate these hybridized artifacts, we disabled secondary alignments by removing the “-a” option in BWA-MEM, forcing each read to align to only its best-matching reference. Given the relatively short length of g*TR* reference sequences, many paired-end reads cannot map as proper pairs within the same reference, potentially leading to the exclusion of informative single-end reads if paired-end criteria are strictly enforced. To more effectively leverage information from both single-end and paired-end reads, we developed an edit distance-based weighting system that assigns differential weights based on read type and mapping characteristics. This system requires perfect matches for reads mapping to core regions while allowing tolerance for variants in flanking regions, reflecting the fact that flanking sequences in our expanded database are derived from merged sequences and may legitimately contain variants. Single-end and paired-end reads receive different tolerance levels and weights within this framework, with paired-end reads generally receiving higher confidence scores due to their increased reliability.

We defined allele depth using customized depth-defining regions (DDRs) that calculate the minimum weighted read depth across specified positions within each allele sequence (**Figure 1**). For most alleles, DDRs are restricted to core regions to ensure both adequate read depth and coverage specificity. However, for nested-structured alleles, where the core sequence of a shorter allele is identical to an internal region of a longer allele (**Supplementary Table 2**), we extended DDRs to include flanking regions to enable proper discrimination these two alleles. The depth calculation at each position incorporates our distance-based weighting system, accounting for read length, reference length, read pairing status, and variant location relative to core and flanking regions.

For alleles exhibiting hybridized structures, we implemented an additional validation to prevent false positive calls resulting from misaligned reads that could artificially combine variants from different true alleles. We clustered all reference alleles based on sequence similarity using Clustal Omega (version 2.3.0) and performed multiple sequence alignments within each cluster (Madeira et al., 2024). Within these alignments, we manually identified key variants as positions that distinguish between alleles within the same cluster, encompassing both allele-dependent variants that differentiate alleles of the same gene level and novel variants that distinguish from their closest IMGT-documented counterparts. For alleles containing multiple key variants, we required that each key variant be supported by at least one properly paired read group where both reads align primarily to the same reference allele, ensuring that the observed variant combination reflects genuine allelic structure rather than alignment artifacts.

### 2. Benchmarking demonstrates high accuracy of the gAIRR-wgs pipeline

The gAIRR-wgs pipeline was benchmarked against 44 HPRC subjects with validated *TR* annotations from personal genome assemblies. When the gAIRR-panel, originally designed for targeted sequencing, was directly applied to WGS data, it achieved suboptimal performance with only 72.5% recall for TR_D and 75.4% recall for TR_V genes, demonstrating the limitations of applying panel-optimized algorithms to WGS data (**Table 1**, **Supplementary Table 3**). In contrast, the optimized gAIRR-wgs pipeline achieved excellent performance with 100% recall and 100% precision for TR_D genes, 99.9% recall and 99.6% precision for TR_J genes, and 99.2% recall and 97.4% precision for TR_V genes. When examining the performance by allele type, the pipeline showed distinct patterns for known versus novel alleles. For documented alleles present in the IMGT database (Version 3.1.41; accessed on 2025-04-19), gAIRR-wgs achieved 100.0% recall and 99.6% precision for TR_J genes, and 99.7% recall and 98.1% precision for TR_V genes. For novel alleles not previously catalogued in reference databases, the pipeline maintained robust performance with 98.5% recall and 97.5% precision for TR_J genes, and 98.3% recall and 93.9% precision for TR_V genes. Notably, these results exceeded the performance of the gAIRR-panel on high-depth targeted sequencing data, demonstrating the effectiveness of our WGS-specific optimizations.

**Table 1.**
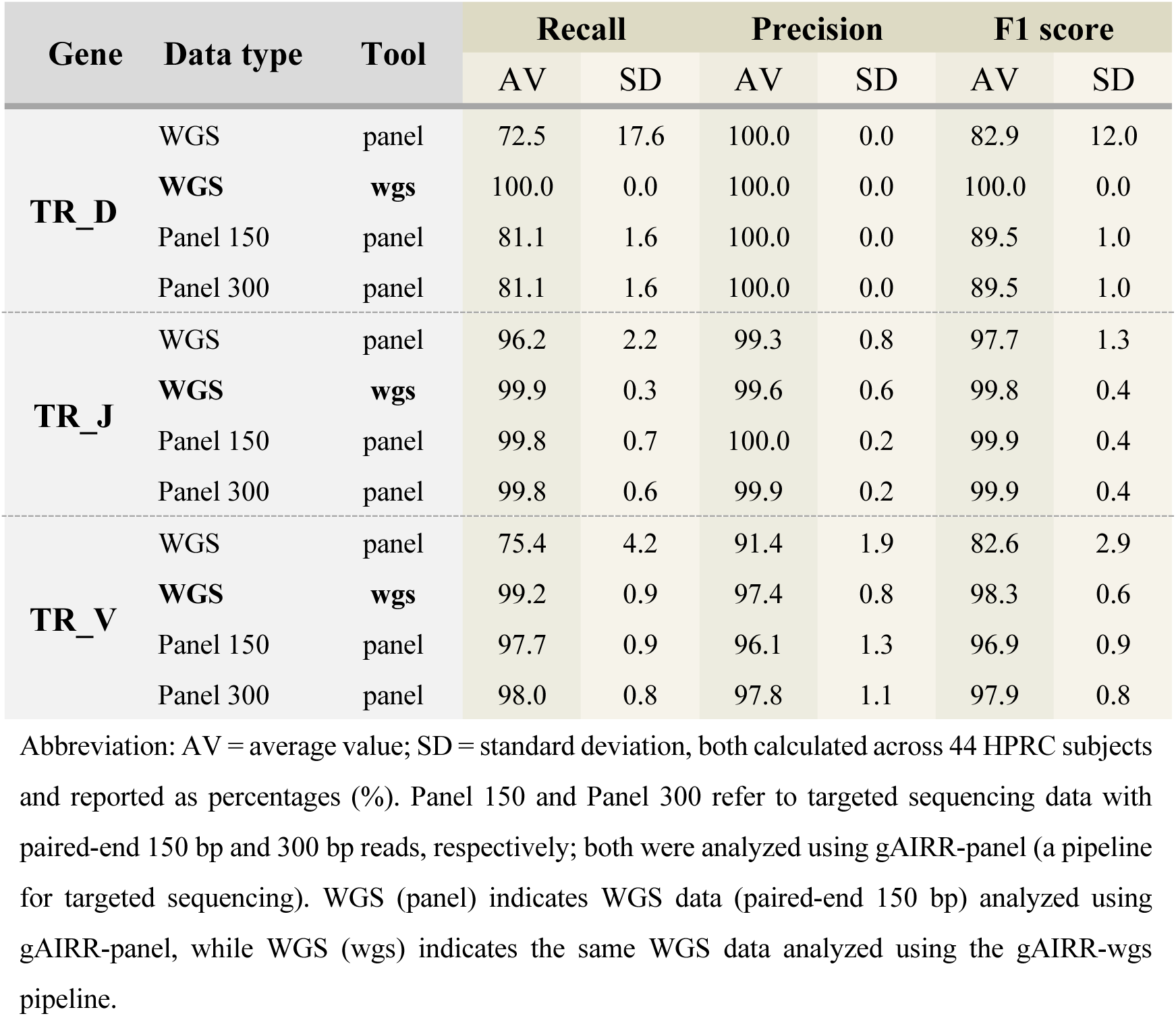
Performance comparison of gAIRR-wgs and gAIRR-panel pipelines across different sequencing platforms in 44 HPRC subjects.

| Gene | Data type | Tool | Recall |  | Precision |  | F1 score |  |
| --- | --- | --- | --- | --- | --- | --- | --- | --- |
|  |  |  | AV | SD | AV | SD | AV | SD |
| TR_D | WGS | panel | 72.5 | 17.6 | 100.0 | 0.0 | 82.9 | 12.0 |
|  | WGS | wgs | 100.0 | 0.0 | 100.0 | 0.0 | 100.0 | 0.0 |
|  | Panel 150 | panel | 81.1 | 1.6 | 100.0 | 0.0 | 89.5 | 1.0 |
|  | Panel 300 | panel | 81.1 | 1.6 | 100.0 | 0.0 | 89.5 | 1.0 |
| TR_J | WGS | panel | 96.2 | 2.2 | 99.3 | 0.8 | 97.7 | 1.3 |
|  | WGS | wgs | 99.9 | 0.3 | 99.6 | 0.6 | 99.8 | 0.4 |
|  | Panel 150 | panel | 99.8 | 0.7 | 100.0 | 0.2 | 99.9 | 0.4 |
|  | Panel 300 | panel | 99.8 | 0.6 | 99.9 | 0.2 | 99.9 | 0.4 |
| TR_V | WGS | panel | 75.4 | 4.2 | 91.4 | 1.9 | 82.6 | 2.9 |
|  | WGS | wgs | 99.2 | 0.9 | 97.4 | 0.8 | 98.3 | 0.6 |
|  | Panel 150 | panel | 97.7 | 0.9 | 96.1 | 1.3 | 96.9 | 0.9 |
|  | Panel 300 | panel | 98.0 | 0.8 | 97.8 | 1.1 | 97.9 | 0.8 |

Abbreviation: AV = average value; SD = standard deviation, both calculated across 44 HPRC subjects and reported as percentages (%). Panel 150 and Panel 300 refer to targeted sequencing data with paired-end 150 bp and 300 bp reads, respectively; both were analyzed using gAIRR-panel (a pipeline for targeted sequencing). WGS (panel) indicates WGS data (paired-end 150 bp) analyzed using gAIRR-panel, while WGS (wgs) indicates the same WGS data analyzed using the gAIRR-wgs pipeline.

To further assess the robustness of our pipeline under conditions that mimic real-world scenarios with incomplete g*TR* databases, we performed additional validation by randomly removing 10%, 30%, 50%, 70%, and even all of the TR_V and TR_J novel alleles from the reference database according to each sample’s annotation results. For TR_J genes, performance remained relatively stable, with both accuracy and sensitivity preserved under 30% removal and high precision still retained even with 50% loss (**Supplementary Figure 4**, **Supplementary Table 4**). In contrast, the more complex and diverse TR_V genes were more vulnerable to database incompleteness: recall dropped substantially once more than 10% of novel alleles were removed, though precision remained relatively high. These findings confirm that our strategy of incorporating previously discovered novel alleles into the database is essential to achieving robust allele calling, and also highlight that incomplete databases can critically constrain performance, particularly for highly diverse gene families such as TR_V. Importantly, the results also demonstrate that once the reference database reaches a sufficient level of completeness, recall of novel alleles can be substantially improved. In other words, a comprehensive database is not only crucial for maintaining accuracy but also indispensable for maximizing sensitivity in novel allele detection.

For comparative evaluation of gAIRR-wgs, we accessed it against ImmunoTyper2, a recently developed state-of-the-art tool for *TR* gene analysis from WGS data (Ford et al., 2025). Both pipelines were applied to the same 44 HPRC subjects used in our validation study. The computational efficiency analysis revealed substantial differences between the two approaches. gAIRR-wgs completed the analysis of all 44 samples in about 6 hours (06:42:50), while ImmunoTyper2 required approximate 7 days (7-19:51:01) to process the same dataset. This represents approximately a 28-fold improvement in computational speed. For known alleles documented in the IMGT database, both tools demonstrated excellent recall performance (**Supplementary Figure 5**). However, gAIRR-wgs achieved superior precision compared to ImmunoTyper2, resulting in fewer false-positive calls and more reliable allele annotations. The combination of high computational efficiency and enhanced precision makes gAIRR-wgs particularly well-suited for biobank-scale analyses and population immunogenomics research, where both processing speed and accuracy are essential for large-scale genomic studies.

### 3. gAIRR-wgs reveals the comprehensive *TR* allele landscape of the Taiwan Han Chinese population Using Taiwan Biobank WGS data

We applied the gAIRR-wgs pipeline to analyze *TR* alleles from 1,492 individuals in the Taiwan Biobank (TWB), revealing a comprehensive picture of *TR* allele diversity in the Han Chinese population. Our analysis identified numerous novel alleles not present in the IMGT database, all supported by high-quality sequencing reads (MAPQ ≥20, AS ≥150) (**Figure 2**, **Supplementary Figure 6**). All novel alleles were systematically renamed using the IgLabel tool to ensure standardized nomenclature (Lees et al., 2023). On average, each individual carried 185.9 (±9.4) TR_V allele types, including 34.6 (±5.5) novel TR_V alleles, 88.5 (±1.9) TR_J allele types with 2.6 (±1.2) novel TR_J alleles, and either 5 (64.1%) or 6 (35.9%) TR_D alleles with very few novel variants observed (**Table 2**, **Supplementary Table 6**). Notably, 109 novel TR_V alleles and 15 novel TR_J alleles exhibited cohort frequencies exceeding 1% in the population, indicating their substantial prevalence rather than rare variants. Considering each allele’s carrier rate in the population, novel alleles accounted for 18.39% of the total TR_V repertoire and 2.77% of the total TR_J repertoire among alleles with cohort frequencies greater than 1% in the TWB cohort.

**Figure 2.**
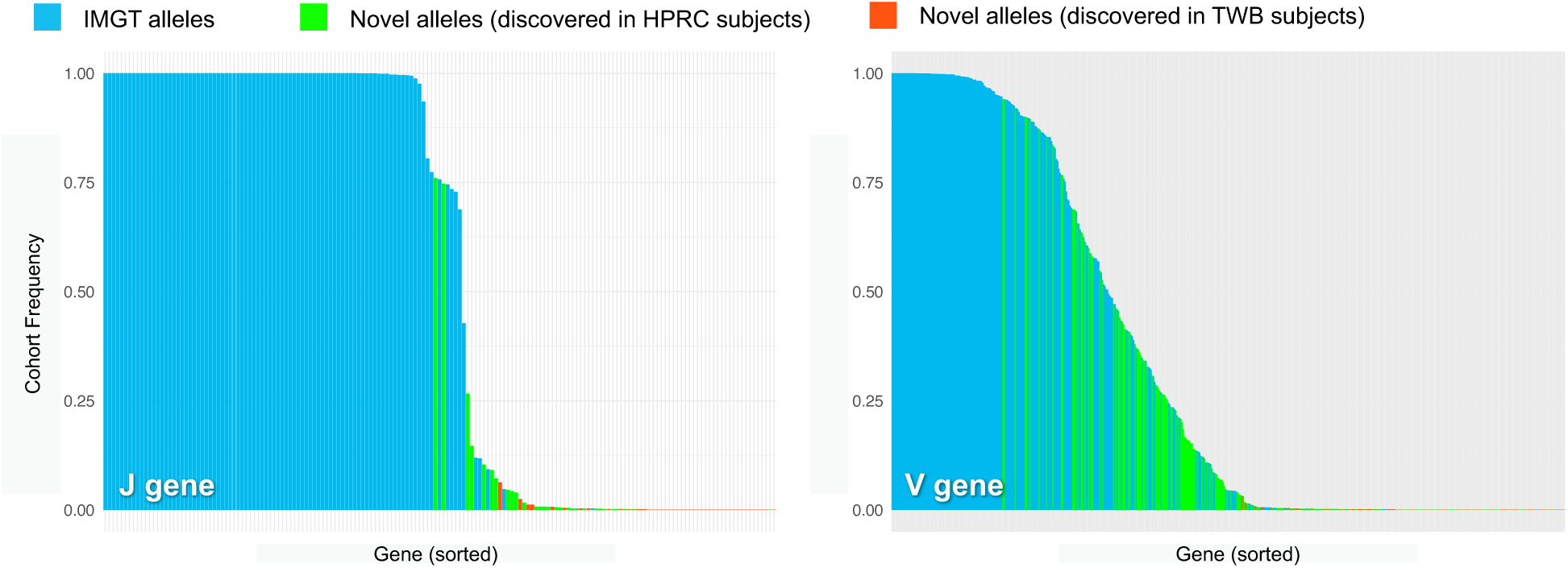
Cohort frequency distribution of TR_J and TR_V alleles in the Taiwan Biobank. Cohort frequency distributions of TR_J (left) and TR_V (right) alleles identified in Taiwan Biobank (TWB) subjects. The Y-axis represents the cohort frequency, defined as the proportion of individuals carrying each specific allele. The X-axis lists individual alleles, sorted in descending order by cohort frequency. Bars are color-coded based on allele type and discovery origin: Blue bars denote alleles already documented in the IMGT database. Green bars represent novel alleles previously discovered in HPRC subjects. Red bars indicate novel alleles discovered only in the TWB and not observed in HPRC.

**Table 2.**
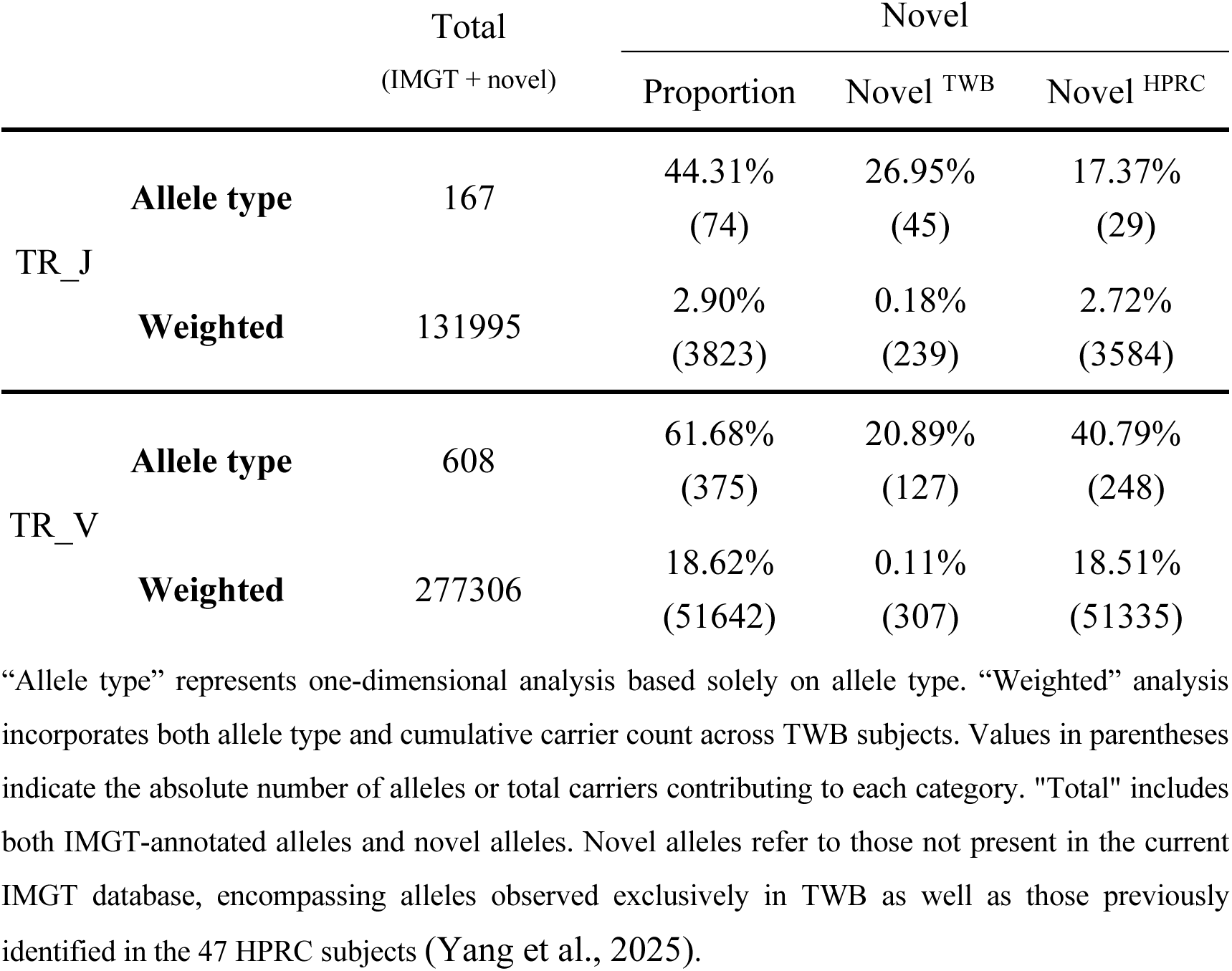
Summary of TR_V and TR_J alleles identified in the TWB cohort.

Given the substantial number of novel alleles discovered, we sought to validate these findings using independent analytical approaches. We primarily employed IGV visualization to manually inspect the sequencing reads, confirming that perfectly matched reads covered the core sequences of these novel alleles and extended into the flanking regions, providing strong evidence for their presence in the samples. Additionally, we cross-validated our findings by examining whether these novel alleles could also be detected using DRAGEN variant calling. We converted all discovered novel alleles to variant format relative to GRCh38 and compared them with a DRAGEN Iterative gVCF Genotyper variant callset from the same TWB samples (as described in the **METHODs** section). Remarkably, many novel alleles were not detected in the DRAGEN callset (**Supplementary Figure 7**, **Supplementary Table 7**), suggesting that conventional variant calling approaches may face challenges in capturing variation within complex genomic regions such as the *TR* loci. While DRAGEN incorporates specialized algorithms for handling nine complex genes including *HLA* and *CYP2D6*, *TR* genes are not included in this targeted optimization, which may contribute to the observed limitations in detecting *TR*-specific variants (Behera et al., 2025). This highlights the necessity for specialized approaches like gAIRR-wgs that are specifically designed to address the unique challenges posed by the highly repetitive and structurally complex nature of *TR* loci.

To gain deeper insights into the diversification of *TR* genes in Taiwan Han Chinese population, we categorized all *TR* genes by cohort frequency into six groups: singleton, doubleton to <0.5%, 0.5% to <1%, 1% to <10%, 10% to <50%, and 50% to 100% (**Figure 3**, **Supplementary Figure 8**). For each *TR* gene, we identified its dominant allele within each cohort frequency group and categorized it as either a IMGT-documented allele or a novel allele (with suffix “_novel”). This allowed us to determine the proportion of genes in each frequency group that were dominated by novel versus known alleles. In the highest frequency group (50-100%), TR_V genes were predominantly represented by *01 alleles (the earliest documented IMGT variants), followed by novel alleles, and then *02 alleles. However, in high-frequency TR_V genes (10-50%), 42.4% were dominated by novel alleles, with *01 and *02 each representing approximately 20%. This pattern indicated substantial database incompleteness, as many genes with greater than 1% cohort frequency were dominated by novel rather than IMGT-documented alleles.

**Figure 3.**
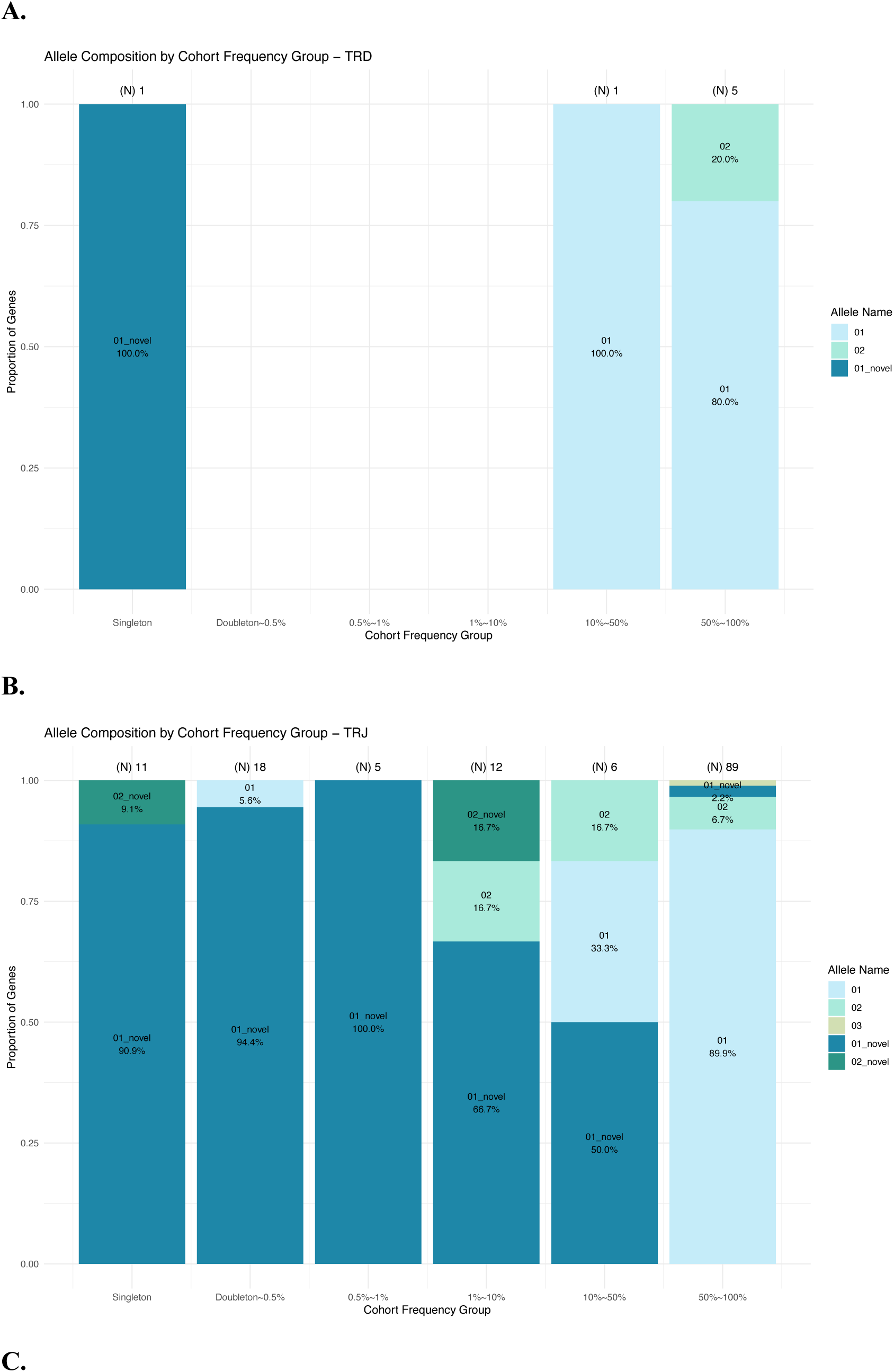
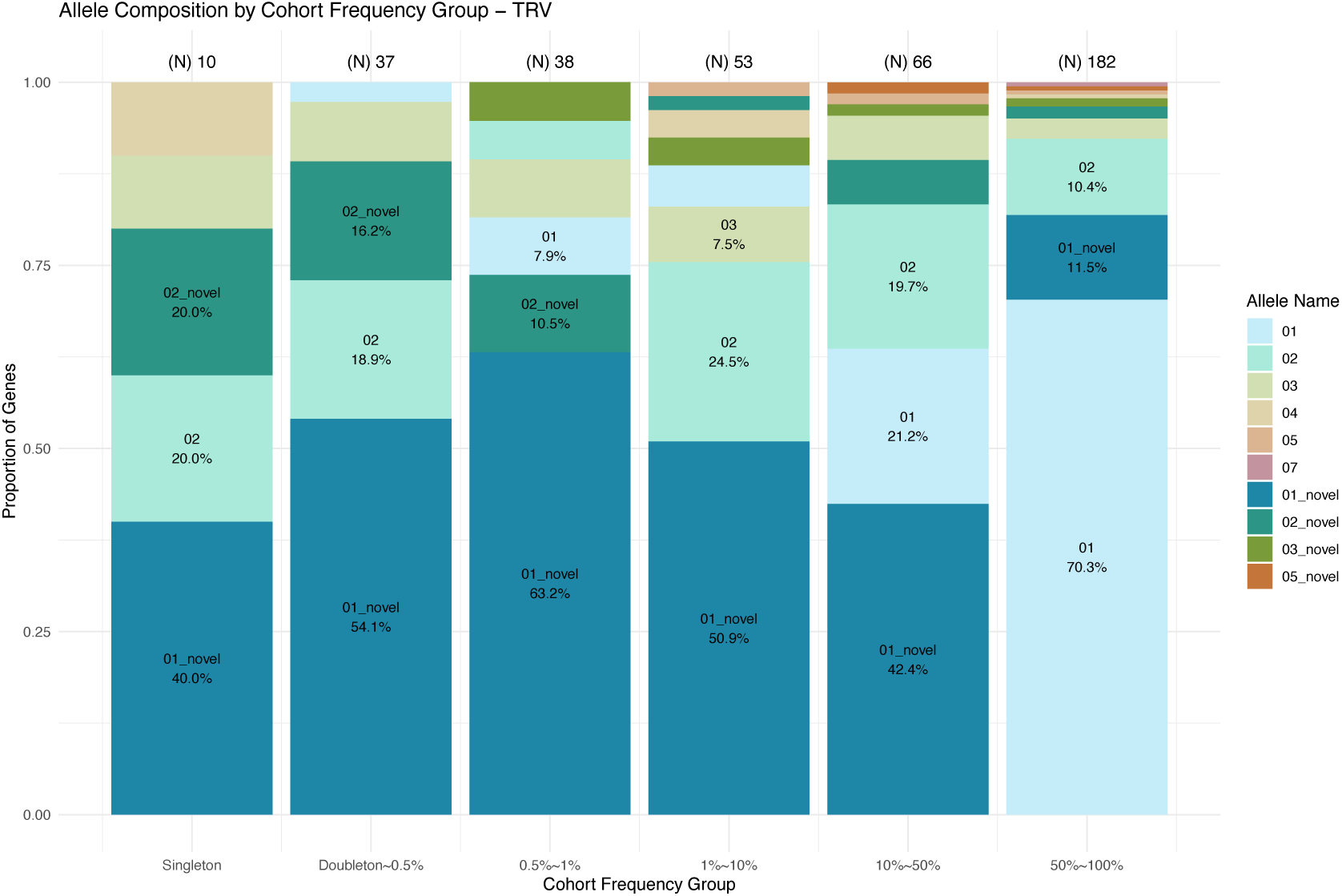
Distribution of Star Alleles Across *TR* Gene Segments Grouped by Cohort Frequency. (A), (B), and (C) show TR_D, TR_J, and TR_V genes, respectively. Proportional distributions of dominant star alleles across *TR* gene segments in the TWB cohort, grouped by cohort frequency. Each gene was assigned to one of six frequency intervals as shown along the X-axis. The Y-axis represented the proportion of genes within each frequency category that were dominated by a given star allele type. Each bar was colored by star allele type, with allele assignments based on either direct IMGT annotation or novel allele grouping (e.g., *01_novel was used to represent novel alleles most similar to the canonical *01). These plots illustrated how allelic composition shifted across frequency categories, revealing evolutionary patterns of conservation and diversification in different *TR* segments. While high-frequency groups were dominated by conserved alleles such as *01, lower-frequency groups increasingly featured novel alleles, particularly *01_novel, reflecting ongoing diversification and potential adaptive emergence within the population.

Among common alleles, with cohort frequencies greater than 1%, 34.17% of TR_V genes (109 out of 319) and 14.02% of TR_J genes (15 out of 107) were dominated by novel alleles (**Figure 4**A, **Supplementary Table 6**). The majority of these high-frequency novel alleles had been previously identified in the 47 HPRC subjects from the first year release data, which were analyzed using long-read sequencing with manual validation (Yang et al., 2025), representing diverse global ancestries and suggesting they represent broader human *TR* diversity missing from current databases. However, a notable proportion appeared to be TWB-specific based on initial comparison: 1.83% of common novel TR_V alleles (2 out of 109) and 26.67% of high-frequency novel TR_J alleles (4 out of 15) were not observed in the initial 47 HPRC subjects. To further investigate this apparent population specificity, we extended our analysis to include 232 HPRC subjects from the HPRC second year data release, which were annotated using gAIRR-annotate (**Figure 4**B). This expanded comparison revealed that most of these previously “TWB-specific” common novel alleles were indeed present in East Asian (EAS) and South Asian (SAS) populations within the larger HPRC cohort (**Figure 4**C, **Supplementary Table 9**). This finding suggests that the initial appearance of population-specific alleles was largely due to limited sample size in the first HPRC release rather than true population restriction. Nevertheless, the broader pattern still highlights the historical underrepresentation of East Asian populations in existing genomic resources and emphasizes the importance of diverse population sampling for comprehensive characterization of *TR* diversity.

**Figure 4.**
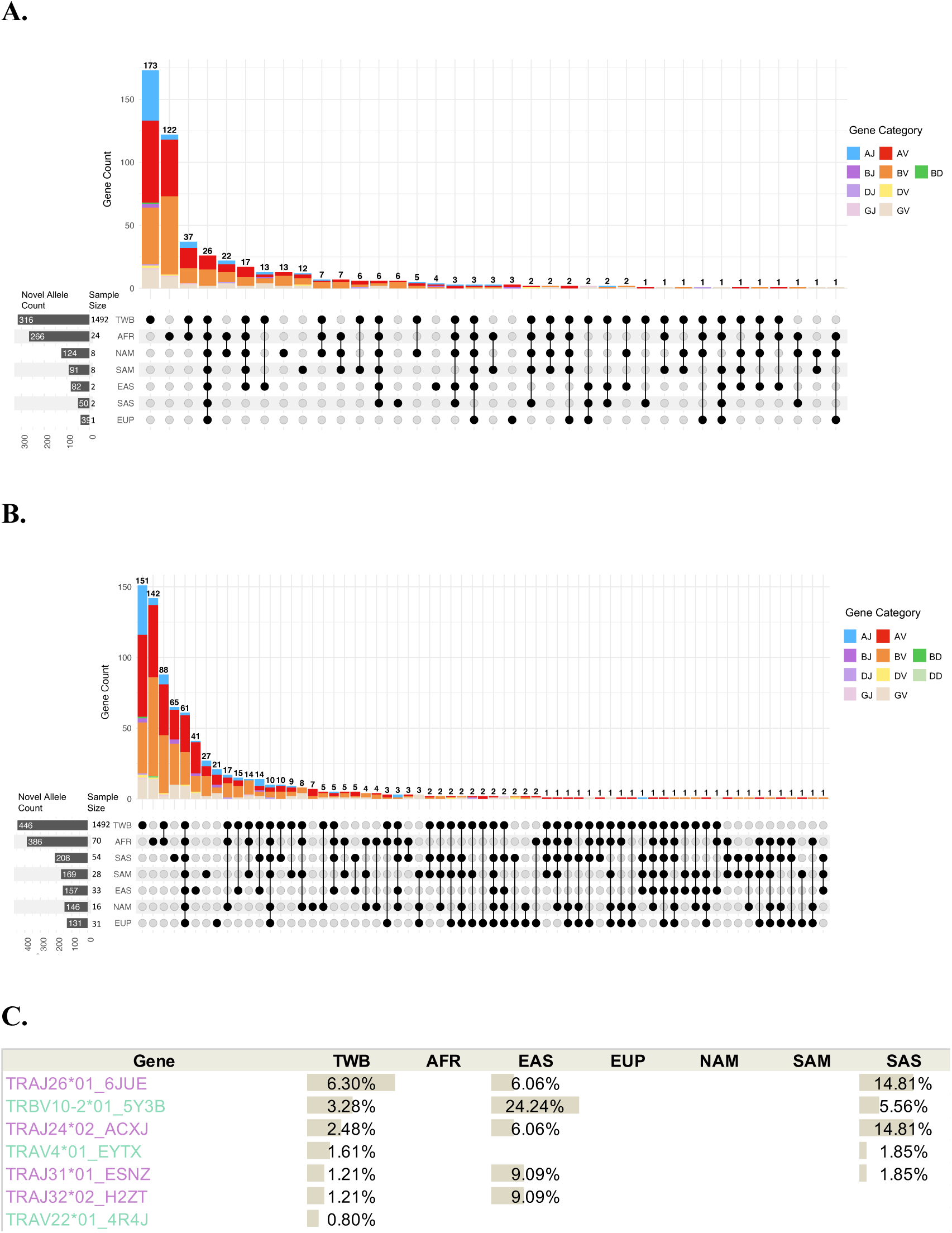
Distribution of novel *TR* alleles across populations. Novel alleles were identified from the Taiwan Biobank (TWB) cohort using the gAIRR-wgs genotyping pipeline. Each bar in the upper panel represents the number of genes carrying novel alleles observed across specific population combinations, with colors indicating gene categories (e.g., AV, AJ, BV, etc.). The lower panel is an UpSet plot summarizing population sharing patterns for these novel alleles across cohorts including TWB, AFR (African), NAM (North American), SAM (South American), EAS (East Asian), SAS (South Asian), and EUP (European). The left margin indicates the total number of novel alleles and sample size per cohort. (A) Novel alleles in TWB were compared against 47 HPRC subjects based on high-confidence annotations from the first HPRC assembly release, using the gAIRR-annotate pipeline with manual validation by long-read alignments. (B) Novel alleles in TWB were compared against 232 HPRC subjects from the second HPRC assembly release, annotated by gAIRR-annotate without long-read-based manual validation. (C) Common novel alleles (>1% frequency) initially appearing to be TWB-specific in the first HPRC release were identified in East Asian (EAS) and South Asian (SAS) populations within the larger set of 232 HPRC subjects from the second release.

High-frequency novel alleles (≥1%) shared across populations likely reflect systematic gaps in reference databases, while the prevalence of TWB-specific alleles across frequency ranges underscores the underrepresentation of ethnic diversity in current immunogenomic databases. Validation against the most recent release of 232 HPRC subjects further corroborated these findings, confirming both the systematic database gaps and insufficient ethnic diversity representation (**Figure 4**).

### 4. Identification of structural deletion polymorphisms in *TR* gene loci

Analysis of *TR* gene presence across the 1,492 TWB subjects using gAIRR-wgs revealed that four genes showed missing rates >2%: *TRGV5* (2.95% missing), *TRGV4* (5.03% missing), *TRBV3-2* (37.13% missing), and *TRBV4-3* (38.50% missing) (**Supplementary Figure 8**). These substantial missing rates suggested the presence of structural deletion polymorphisms rather than technical artifacts, prompting further investigation into the genomic architecture of these regions.

The affected genes exhibited distinct chromosomal arrangements. *TRGV4* and *TRGV5* are physically adjacent on chromosome 7 (GRCh38), separated by approximately 4.1 kb, while *TRBV3-2* and *TRBV4-3* are located on the alternative contig chr7_KI270803v1_alt, separated by approximately 3.1 kb. To validate these deletion events, we examined alignment data from TWB WGS samples processed using the GATK best practices pipeline with BWA-MEM mapping. Visual inspection revealed complete absence of read support at the genomic positions of these genes in affected individuals. Furthermore, we observed coverage dropouts spanning approximately 9 kb encompassing *TRGV4* and *TRGV5*, and approximately 21 kb encompassing *TRBV3-2* and *TRBV4-3* (**Figure 5**, **Supplementary Figure 12**). Identical patterns were observed in alignment data generated using DRAGMAP, ruling out mapper-specific artifacts and supporting the hypothesis of genuine deletion events.

**Figure 5.**
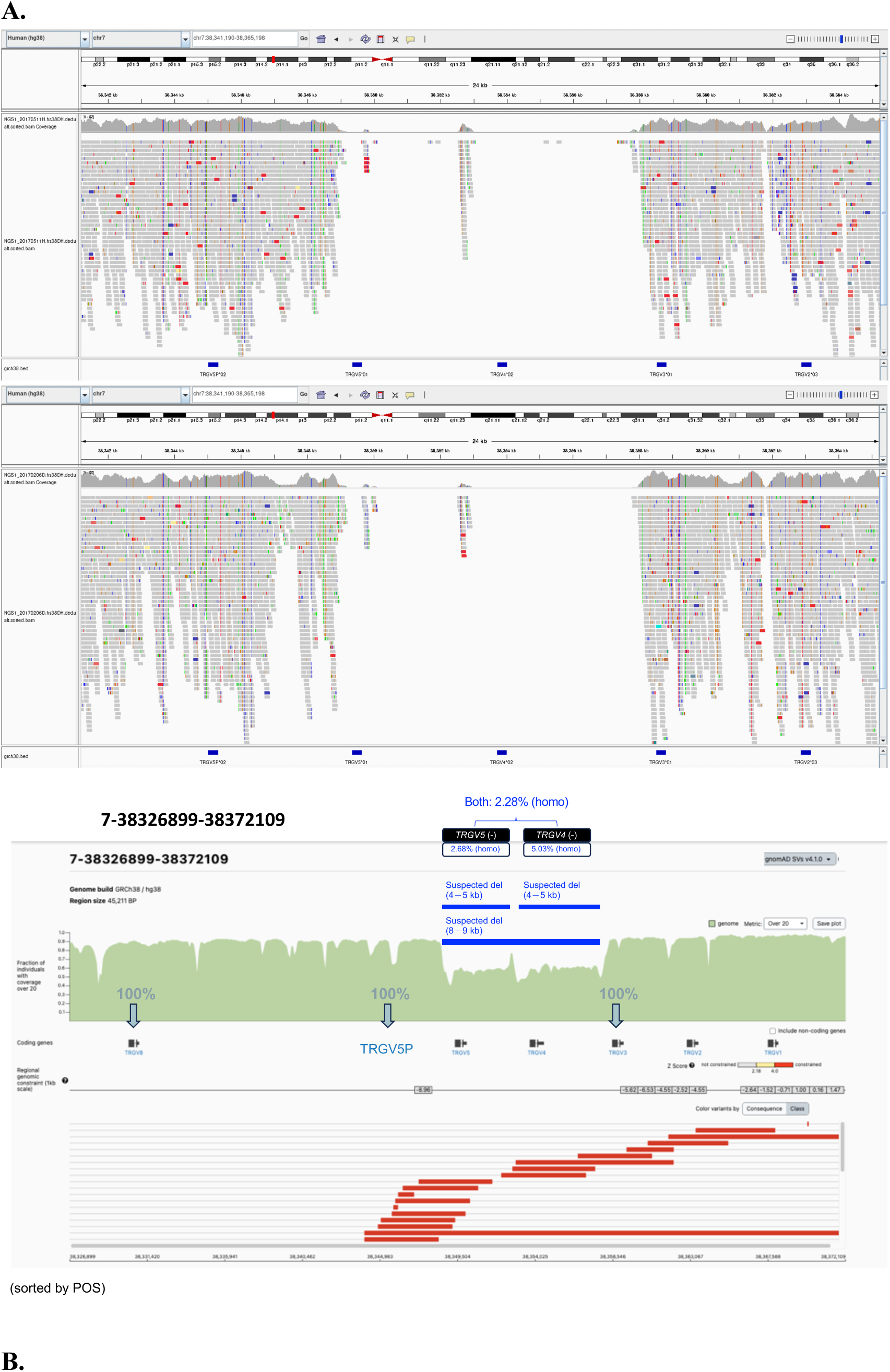
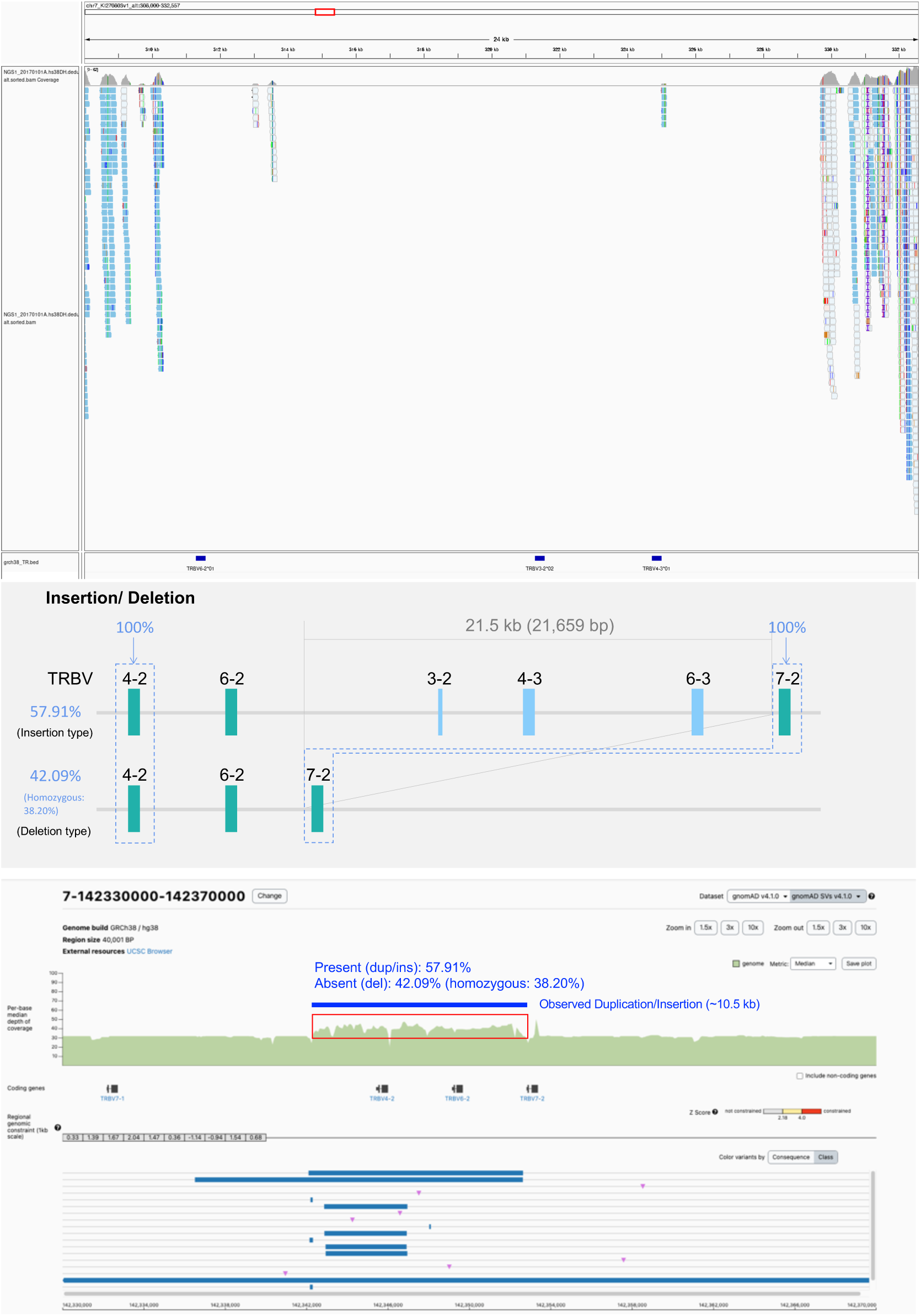
IGV visualization of structural variation at *TRG* and *TRB* loci in representative TWB WGS samples. (A) T*RGV4*/*TRGV5* deletion polymorphism. Read alignments from GATK-processed WGS data mapped to GRCh38 are shown. Affected individuals exhibit complete absence of read coverage across the *TRGV4* and *TRGV5* region, with sharp dropouts in depth, indicating a homozygous deletion. Genes *TRGV4* and *TRGV5* are physically adjacent on chr7, separated by ∼4.1 kb. Below, comparison with SV databases highlights the expected deletion interval (blue box), with TWB carrier frequencies indicated (Present: 97.7%, Absent: 2.28%). (B) *TRBV3-2*/*TRBV4-3* insertion/deletion polymorphism. In a representative deletion haplotype, no read coverage is observed at the positions of *TRBV3-2* and *TRBV4-3* (∼21 kb span), while neighboring genes *TRBV4-2* and *TRBV7-2* retain full coverage, consistent with their stable presence. Read alignment visualizations were generated using IGV (Integrative Genomics Viewer). SV database comparison is shown below, with the predicted deletion span marked in blue and TWB carrier frequencies annotated.

#### *TRGV4*/*TRGV5* Deletion Polymorphism

Our findings corroborate previous reports of genomic deletions in this region. One study documented homozygous genomic deletion encompassing *TRGV4* and *TRGV5* in 1 individual of South Asian ancestry within a cohort of 45 subjects (Corcoran et al., 2023). Using gAIRR-wgs, we identified a homozygous carrier rate of 2.28% for the absence of both genes on two haplotypes in the TWB cohort, suggesting that the event may arise from a ∼9 kb deletion encompassing both genes, as inferred from IGV inspection (**Figure 5**). We compared our findings with the gnomAD SVs database (v4.1.0) and observed notable discrepancies. Only six smaller deletions (4∼5 kb) were identified in gnomAD, involving *TRGV4* and *TRGV5* separately. The larger ∼9 kb deletion encompassing both genes that we observed was not reported. This discrepancy likely reflects the fact that the *TR* region may not fall within gnomAD’s high-confidence callable regions. Indeed, inspection of the gnomAD database revealed that approximately half of the samples showed poor sequencing quality in this region, and although several deletion affecting *TRGV4* and *TRGV5* were detected, they may have been filtered out during quality control procedures, potentially leading to underestimation of the true deletion frequency in this region.

#### *TRBV3-2*/*TRBV4-3* Insertion/Deletion Polymorphism

The absence patterns of *TRBV3-2* and *TRBV4-3* align with previously described structural variation in the *TRBV* locus. Literature reports and IMGT documentation describe a 21.5 kb insertion/deletion polymorphism encompassing *TRBV3-2*, *TRBV4-3*, and *TRBV6-3* (Omer et al., 2022; Zhao et al., 1994). Although *TRBV6-3* was excluded from our gAIRR-wgs analysis due to its identical sequence to upstream *TRBV6-2*, making them indistinguishable, we were able to characterize this polymorphism using the presence/absence patterns of *TRBV3-2* and *TRBV4-3* as proxy markers. Our analysis revealed that 38.20% of TWB individuals carried the homozygous deletion haplotype (absence of both *TRBV3-2* and *TRBV4-3* on both haplotypes), while 57.91% carried the insertion haplotype (presence of both genes on at least one haplotype). Furthermore, analysis of the 232 assemblies from the HPRC release 2 dataset revealed that 34.48% of individuals carried the homozygous deletion type, with the highest proportion observed in AFR (12.93% of 70 individuals), SAS (9.48% of 54 individuals) and EAS (3.45% of 33 individuals) (**Supplementary Table 10**). Due to the limitations of gAIRR-wgs, only the overall carrier rate of the insertion haplotype could be estimated (57.91%); when applying the same perspective to the 232 HPRC assemblies, we observed a comparable carrier rate of approximately 50%. Furthermore, we searched gnomAD SVs (v4.1.0) for corresponding structural variants (**Figure 5**). Since the GRCh38 primary contig represents the deletion haplotype, we queried for duplications/insertions between *TRBV4-2* (chr7:142345421-142345985) and *TRBV7-2* (chr7:142352819-142353358). One duplication showed overlap with our observed insertion haplotype: a 10,548 bp duplication (DUP_CHR7_C54B5731) with ∼54.72% carrier frequency in East Asians. However, the expected ∼21 kb insertion originates from a historical duplication, resulting in high sequence similarity between *TRBV4-2*, *TRBV6-2*, and their flanking regions. Consequently, short reads derived from this duplicated segment tend to align back to the GRCh38 primary contig, which represents the deletion haplotype lacking *TRBV3-2*, *TRBV4-3*, and *TRBV6-3*. As a result, many reads are preferentially mapped to *TRBV4-2* and *TRBV6-2*, with some alignments lost, leading to the duplication being interpreted as ∼10.5 kb rather than ∼21 kb. When considering the GRCh38 alternative contig (chr7_KI270803v1_alt), which represents the insertion haplotype, reads can be correctly aligned to the intended loci, thereby revealing the presence of the full ∼21 kb structural variant among individuals carrying the insertion type.

In parallel, we compared our observations with structural variants called by the Manta SV caller (DRAGEN pipeline) on the same TWB dataset. No SV segments corresponding to the *TRGV* deletion or the *TRBV* insertion/deletion polymorphism were identified (**Supplementary Table 12**, **Supplementary Table 11**). This discrepancy underscores the inherent difficulty of resolving complex immune gene regions using conventional SV callers.

### 5. Population-Specific *TR* Allele Distribution Patterns and Diversity Analysis

To systematically characterize population-specific patterns of *TR* allele diversity, we analyzed allele distributions across the TWB and 232 HPRC subjects representing six global populations: African (AFR, n=70), North American (NAM, n=16), South American (SAM, n=28), East Asian (EAS, n=33), South Asian (SAS, n=54), and European (EUP, n=31) (**Supplementary Figure 9**, **Supplementary Table 8, Supplementary Table 9**).

We employed Hellinger distance calculations to quantify differences in allele frequency distributions between populations, with a threshold of >0.5 indicating significant distributional differences (refer to **METHODS**). Overall, 57 *TR* genes (56 TR_V and 1 TR_J) showed significant distributional differences (Hellinger distance >0.5) in at least one population comparison (**Supplementary Figure 9**, **Supplementary Table 13**). The African population (AFR) showed the most distinctiveness, with 50 genes exhibiting significant differences when comparing AFR against each of the other six populations, totaling 151 gene-population pair comparisons with Hellinger distance >0.5. On average, each significantly different gene in AFR differed from 3.02 other populations. The East Asian population (EAS) demonstrated the second-highest level of differentiation with 44 genes showing significant differences across 60 gene-population comparisons, averaging 1.36 population differences per gene. The TWB exhibited 34 genes with significant differences, generating 78 gene-population pair comparisons where TWB differed from other populations, averaging 2.29 population differences per gene. South American (SAM) and South Asian (SAS) populations showed the least differentiation, with 28 genes (36 comparisons) and 30 genes (33 comparisons) respectively.

Among the TWB cohort, 34 *TR* genes (33 TR_V and 1 TR_J) exhibited significantly different allele distributions compared to other global populations (Hellinger distance >0.5). Seven TR_V genes showed particularly pronounced differences: *TRBV20-1*, *TRAV6*, *TRAV35*, *TRBV7-5*, *TRAV12*-2, *TRAV8-4*, and *TRBV5-5*. When comparing TWB specifically with the EAS population, most *TR* genes showed similar allele distributions, reflecting shared ancestry. However, eight TR_V genes maintained significant distributional differences even within this regional comparison, including the seven genes mentioned above plus *TRAV38-1*, suggesting population-specific evolutionary pressures or founder effects within East Asian subpopulations.

Moreover, we calculated Shannon’s diversity index for each gene locus across all cohorts to quantify allelic richness within populations (see **METHODS**). Using the first quartile (Q1) value of 0.4793 across all results as a threshold, we defined high diversity genes, identifying 104 *TR* genes spanning the seven cohorts that exceeded this threshold (**Supplementary Figure 11**, **Supplementary Table 14**). AFR populations exhibited the highest overall diversity, with 84 genes meeting the high diversity criterion, consistent with established patterns of greater genetic diversity in African populations. In contrast, TWB showed the lowest diversity with only 47 high-diversity genes.

Notably, 25 *TR* genes (24 TR_V and 1 TR_J) maintained consistently high diversity across all seven cohorts, representing core genes with universally preserved allelic richness. Conversely, six TR_V genes (*TRBV30*, *TRAV27*, *TRBV20/OR9-2*, *TRBV6-6*, *TRGV5P*, *TRBV5-5*, *TRBV7*, *TRBV25/OR9-2*, *TRAV26-1*, *TRBV7-7*) showed high diversity in other populations but reduced diversity in TWB (Shannon’s index <0.4793), potentially indicating population-specific bottlenecks or selective pressures affecting these loci.

Notably, consistent with our earlier findings regarding limited allelic variation in TR_D segments, diversity analysis revealed that most TR_D genes were dominated by single prevalent alleles across populations (**Supplementary Figure 8**). However, *TRBD2* emerged as a notable exception, exhibiting the highest Shannon’s diversity index among all TR_D genes across different populations. This pattern suggests that while most TR_D segments are highly conserved, *TRBD2* maintains functional diversity that may be important for T cell receptors functionality.

## DISCUSSION

This study addresses a critical gap in immunogenomics by developing the specialized pipeline for accurate *TR* allele calling from standard WGS data. The challenges inherent in applying existing *TR* genotyping tools to WGS data, including reduced sequencing depth, shorter read lengths, and limited paired-end information utilization, have prevented large-scale population studies of g*TR* diversity. Our gAIRR-wgs pipeline overcomes these fundamental limitations through three key innovations: targeted read extraction that reduces computational burden, enhanced reference databases incorporating population-specific diversity, and optimized algorithms that prevent hybridized artifacts while maintaining high sensitivity.

The improvement in computational efficiency, from 6-7 hours and 500GB memory to 12-15 minutes and 5GB memory per sample, represents an improvement that makes population-scale *TR* genotyping feasible for biobank-sized cohorts. This efficiency gain is crucial for translating immunogenomic discoveries to clinical applications, where rapid and cost-effective analysis is essential. The superior performance of gAIRR-wgs compared to gAIRR-call demonstrates that WGS-specific optimizations are not merely adaptations but fundamental improvements in algorithmic design.

To demonstrate the practical utility of gAIRR-wgs and validate its performance in real-world applications, we applied our pipeline to a large-scale population cohort from the Taiwan Biobank. This analysis of 1,492 individuals not only serves as a proof-of-concept for the scalability and robustness of our approach but also provides an unprecedented opportunity to systematically characterize *TR* allelic diversity in a well-defined Han population in East Asia. The choice of TWB as our demonstration cohort is particularly valuable given the historical underrepresentation of East Asian populations in immunogenomic studies, allowing us to assess both the technical performance of gAIRR-wgs and the completeness of existing reference databases in a previously understudied population context.

Our comprehensive analysis of 1,492 TWB individuals reveals profound gaps in current immunogenetic reference databases, particularly for East Asian populations. The finding that 42.4% of high-frequency TR_V genes are dominated by novel alleles rather than IMGT-documented alleles represents a systematic bias in global immunogenetic resources. This disparity is particularly striking given that many of these novel alleles were subsequently found in HPRC subjects of diverse ancestries, and further validation using the expanded HPRC release 2 dataset confirmed that most high-frequency novel alleles initially appearing TWB-specific were indeed present in Asian populations. This progression from apparent population specificity to broader population presence underscores how limited sampling in reference databases can create imperfect impressions of population uniqueness.

Inaccurate or incomplete g*TR* profiling directly impacts our understanding of immune repertoire biases in disease contexts, potentially leading to misinterpretation of adaptive immune responses in non-European populations. The identification of population-specific high-frequency alleles further emphasizes the critical need for population-specific immunogenomic studies. This is precisely where gAIRR-wgs demonstrates its unique value by enabling systematic characterization of *TR* diversity from existing biobank WGS short read data, our approach can rapidly expand reference databases and correct population-specific biases.

Through our gAIRR-wgs post-analysis, we identified structural polymorphisms affecting contiguous *TR* genes, representing a previously underappreciated source of immune diversity. The *TRBV3-2*/*TRBV4-3* insertion/deletion polymorphism, present in Taiwan Han Chinese individuals, demonstrates that structural variants can dramatically alter the available germline *TR* repertoire at the population level. Similarly, the *TRGV4*/*TRGV5* deletion polymorphism represents another example of how structural variants contribute to immune genetic diversity. These findings have profound implications for understanding population-specific immune capabilities and disease susceptibilities. Notably, the failure of structural variant callers with standard/default pipeline to detect these immunologically relevant polymorphisms, despite their high frequency and substantial genomic span, underscores the necessity of specialized approaches for complex immune gene regions. These structural variations are incomplete in the gnomAD database, highlighting systematic gaps in structural variant documentation for both Asian populations and immune-relevant genomic regions. The gene-centric approach employed by gAIRR-wgs proves essential for identifying functionally relevant structural variants that conventional genome-wide methods miss.

While gAIRR-wgs represents a significant advance in *TR* genotyping from WGS data, several technical and methodological limitations warrant consideration and provide directions for future development. The reliance on short-read sequencing technology (PE 150 bp) limits resolution of the most complex structural variants and haplotype phasing. Hybridized-structured alleles, where different segments share similarity with distinct reference alleles, pose particular challenges that could be better resolved with longer reads (≥300 bp) or long-read approaches. Through our benchmark studies, we have identified specific alleles that frequently generate false positives in 30X WGS PE 150 bp data, and we provide a list of these problematic alleles to assist users in recognizing and mitigating potential sources of error.

Additionally, gAIRR-wgs currently does not incorporate relative genomic positioning information, making it unable to distinguish alleles with identical sequences but different chromosomal locations (e.g., TRBV6-2*01/TRBV6-3*01, TRBV24-1*02/TRBV24/OR9-2*01, and TRGJ1*02/TRGJ2*01). While evolutionary considerations may justify distinct nomenclature based on gene duplication history, the assignment of completely different gene names to functionally identical sequences can create confusion in downstream application and clinical interpretation.

The current implementation focuses on carrier status (presence/absence of genes) rather than providing quantitative assessment of allelic dosage. For population-level studies, carrier status and allelic dosage represent different analytical perspectives: carrier analysis effectively captures population-level allele frequencies and diversity patterns, while dosage assessment provides individual-level genotypic resolution. Future versions should incorporate copy number estimation to provide more comprehensive genotyping information, particularly important for understanding gene expression levels and dosage effects in clinical contexts.

Furthermore, gAIRR-wgs currently restricts novel allele calling to core coding regions, following IMGT nomenclature rules. However, we recognize that future implementations should consider non-coding regions, including flanking sequences, for comprehensive allelic characterization, similar to the nomenclature systems used for other complex immune genes such as *CYP*, *KIR*, and *HLA*. While gAIRR-wgs includes algorithms to handle nested structure alleles, some evidence suggests that certain IMGT reference sequences may contain errors. Future work should incorporate systematic validation of reference sequences and provide frameworks for updating problematic entries.

### Conclusion

The gAIRR-wgs pipeline addresses fundamental challenges in population immunogenetics by enabling accurate *TR* allele calling from standard WGS data. Our analysis reveals extensive previously uncharacterized *TR* diversity in Asian populations, identifies structural polymorphisms, and provides frameworks for optimal database expansion strategies. These advances lay the foundation for population-scale immunogenomic studies and precision approaches to immune-related diseases. The systematic characterization of human *TR* diversity across global populations will be essential for understanding immune system evolution, disease susceptibility, and therapeutic response in our increasingly interconnected world. Most importantly, this work demonstrates that advanced computational approaches can unlock the immunogenomic potential of existing biobank resources, accelerating translation of population genetic insights into clinical applications. As precision medicine increasingly recognizes the central role of immune system diversity in health and disease, tools like gAIRR-wgs will be essential for realizing the promise of personalized immunotherapy and population-specific medical interventions.

## METHODS

### 1. gAIRR-wgs pipeline development

We developed gAIRR-wgs specifically optimized for *TR* genotyping from WGS data. The original gAIRR-Suite, includes a module, gAIRR-call (referred to as gAIRR-panel in this manuscript), was designed for targeted analysis of *IG* and *TR* gene analysis using high-depth panel sequencing data (Lin et al., 2022).

#### Read Extraction

WGS reads were first aligned to the GRCh38 reference genome (hs38DH) using BWA-MEM (v.0.7.17) (Li, 2013). Target reads covering *TR* gene loci were extracted, including both mapped reads from target regions and unmapped reads to account for potential reference genome discrepancies.

#### Reference Database Construction

.The *TR* allele reference database was constructed based on the IMGT database, with *TR* gene sequences last updated in 2020 (Manso et al., 2021). The database was then expanded by incorporating 335 novel *TR* alleles previously identified and validated from 47 HPRC personal genome assemblies (Yang et al., 2025).

#### Allele Calling Algorithm

The gAIRR-wgs pipeline follows a two-step alignment process. Initial BWA-MEM alignment identifies candidate novel alleles, which are then incorporated into the reference database for a second alignment round to prevent hybridized artifact formation.

#### Depth Calculation Algorithm

Allele depth was calculated using the minimum coverage principle across core gene regions, with weighted incorporation of both single-end and paired-end reads. The adaptive threshold from gAIRR-panel, which was designed for high-depth panel data, was removed in gAIRR-wgs as it proved suboptimal for WGS data.

Additional details of the computational implementation and parameter optimization are provided in **Supplementary Note 1**.

### 2. Benchmark study

Benchmark validation used WGS data from 44 HPRC subjects (paired-end 150 bp reads) with validated *TR* allele annotations derived from personal genome assemblies and long read sequencing data (Yang et al., 2025). Performance was evaluated using recall (sensitivity) and precision (positive predictive value) metrics:

Recall = True Positives / (True Positives + False Negatives)

Precision = True Positives / (True Positives + False Positives)

Performance metrics are presented as the mean ± standard deviation across all subjects.

Novel alleles were compared based on core sequence identity, with specific exclusions for allele pairs indistinguishable by short-read sequencing (see **Supplementary Note 2** for details).

### 3. Population study

#### Study Cohort and Data Processing

The population analysis included 1,492 individuals from the Taiwan Biobank with Illumina-based WGS data (paired-end 150 bp reads, sequenced using HiSeq 2500 (n=555), HiSeq 4000 (n=634) and NovaSeq 6000 (n=303)) derived from peripheral blood mononuclear cell (PBMC) DNA samples (Hsu et al., 2024). Raw data were provided in CRAM format and processed using our read extraction pipeline to obtain TR-specific reads prior to gAIRR-wgs analysis.

#### Variant Validation and Database Comparisons

For variant-based validation, all novel alleles were aligned to the GRCh38 reference genome (hs38DH) using BWA-MEM with secondary alignments excluded. Novel allele sequences were converted to variant format relative to GRCh38, processing various CIGAR operations including matches (M), insertions (I), deletions (D), substitutions, and clipping events (S, H) were processed alongside MD tags to accurately identify sequence differences and generate REF to ALT variant calls for comparison with the DRAGEN Iterative gVCF Genotyper variant callset. Variants were validated based on concordance in genomic positions, allele sequences, and genotype calls with adequate coverage depth (detailed validation criteria in **Supplementary Note 3**). For reference blocks (regions marked with <NON_REF> in ALT field representing sequences identical to the reference genome), validation was performed by confirming the absence of variant calls within the corresponding genomic intervals.

Structural variant analysis was conducted using results generated by the DRAGEN pipeline (version 4.0.3) on the TWB WGS dataset. These results are derived on unpublished data, and the related manuscript is currently in preparation. For regions with potential structural variants identified through gene absence patterns, corresponding genomic intervals were systematically queried in the DRAGEN SV callset to identify overlapping or similar-length structural variants. SV calls were considered concordant when: (1) genomic coordinates showed substantial overlap with predicted deletion/insertion regions, and (2) variant sizes were within reasonable similarity ranges of the expected structural changes based on gene absence frequencies and genomic context.

#### Shannon’s Diversity Index Calculation

We calculated Shannon’s diversity index for each *TR* gene locus across all populations to quantify allelic richness and evenness within populations using the formula:

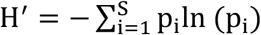

where p_i_ represents the proportion of individuals carrying allele i, and S is the total number of alleles for a given gene. The Shannon’s diversity index ranges from 0 ≤ H′ ≤ ln(S), where H′ = 0 indicates a population dominated by a single allele (no diversity), and higher values indicate greater allelic diversity. We used the first quartile (Q1) value of 0.4793 across all gene-population combinations as a threshold to define high-diversity genes.

#### Hellinger Distance Calculation

To quantify differences in allele frequency distributions between populations for each *TR* gene, we calculated Hellinger distance using the formula:

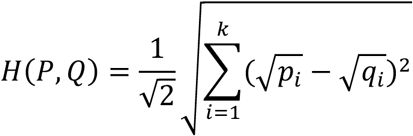

where P and Q represent allele frequency distributions for a given gene in two different populations, p_i_ and q_i_ are the frequencies of allele i in populations P and Q respectively, and k is the total number of unique alleles observed across both populations. The Hellinger distance ranges from 0 to 1, where 0 indicates identical allele distributions between populations and 1 indicates completely different distributions. We used a threshold of H > 0.5 to identify genes with significantly different allele distributions between population pairs.

### 4. Statistical Analysis and Visualization

The core tool, gAIRR-wgs was primarily developed in Python (version 3.7.13). All the downstream analyses, including benchmark study and population study, were performed using Python (version 3.10.14) and R (version 4.4.0).

Read alignment visualization and validation were conducted using the Integrative Genomics Viewer (version 2.10.3; IGV) (Robinson et al., 2011). To ensure reproducibility and maintain environment consistency, all computational tools were managed using conda (version 24.11.3), with separate virtual environments created for each major component of the analysis pipeline.

## DATA ACCESS

Short-read sequencing data for the HPRC individuals (HG002, HG00438, HG005, HG00621, HG00673, HG00733, HG00735, HG00741, HG01071, HG01106, HG01109, HG01175, HG01243, HG01258, HG01358, HG01361, HG01891, HG01928, HG01952, HG01978, HG02055, HG02080, HG02109, HG02145, HG02148, HG02257, HG02572, HG02622, HG02630, HG02717, HG02723, HG02818, HG02886, HG03098, HG03453, HG03486, HG03492, HG03516, HG03540, HG03579, NA18906, NA19240, NA20129, and NA21309) were obtained from the Human Pangenome Reference Consortium (HPRC) project. The corresponding SRA or ENA accession numbers are as follows: HG002 (SRR14724532), HG00438 (ERR3988768), HG005 (SRR14724528), HG00621 (ERR3988803), HG00673 (ERR3988814), HG00733 (ERR3988823), HG00735 (ERR3988824), HG00741 (ERR3988826), HG01071 (ERR3988832), HG01106 (ERR3988841), HG01109 (ERR3988842), HG01175 (ERR3988851), HG01243 (ERR3988858), HG01258 (ERR3988862), HG01358 (ERR3988875), HG01361 (ERR3988876), HG01891 (ERR3988943), HG01928 (ERR3988951), HG01952 (ERR3988958), HG01978 (ERR3988965), HG02055 (ERR3988979), HG02080 (ERR3988986), HG02109 (SRR11124438), HG02145 (ERR3988995), HG02148 (ERR3988996), HG02257 (ERR3989003), HG02572 (ERR3989028), HG02622 (ERR3989038), HG02630 (ERR3989040), HG02717 (ERR3989059), HG02723 (ERR3989060), HG02818 (ERR3989080), HG02886 (ERR3989091), HG03098 (ERR3989119), HG03453 (ERR3989166), HG03486 (ERR3989170), HG03492 (ERR3989173), HG03516 (ERR3989174), HG03540 (ERR3989177), HG03579 (ERR3989180), NA18906 (ERR3989364), NA19240 (ERR3989410), NA20129 (ERR3989456), and NA21309 (SRR10392718).

Annotation results and the discovery of 335 novel alleles are provided in the supplementary materials available at https://www.biorxiv.org/content/10.1101/2025.05.24.655452v1.

Short-read and alignment data for the Taiwan Biobank (TWB) cohort were accessed under ethical approval TWBR11106-05. The GRCh38 BED reference used for targeted *TR* allele analysis is publicly available at https://github.com/maojanlin/gAIRRsuite.

## ACKNOWLEDGMENTS

We thank all the participants from the Taiwan Biobank and all investigators who contributed to this study. We are also grateful to the National Center for High-performance Computing (NCHC) of the National Institutes of Applied Research (NIAR) of Taiwan for providing computational and storage resources. This study was supported by the research grants (NSTC 111-2320-B-002-091-MY3, 112-2622-B-002-012) and industry-academia cooperative research program grants (NSTC 112-2622-B-002-012) from the National Science and Technology Council in Taiwan. This project is also supported by the Excellence in Key Advantages Program (NTU Core Consortiums and Competitiveness Programs) from the National Taiwan University (113L892901).

