## Supplementary Table and notes for "gAIRR-wgs: An Algorithm to Genotype T Cell Receptor Alleles Using Whole Genome Sequencing Data": Supplementary_Note.docx

**Supplementary Note 1. Technical Implementation Details of gAIRR-wgs Pipeline.**

**Read Extraction**

**Reference Database Construction**

**Allele Calling Algorithm**

**Depth Calculation Algorithm**

**Supplementary Note 2. Benchmark Validation Details.**

**Handling of Sequence-Identical Allele Pairs**

**Supplementary Note 3. Detailed Validation Criteria for Novel Allele Variants.**

**CIGAR Operation Processing**

**Variant Matching Criteria**

Validation criteria for novel allele variants against the DRAGEN callset were
